# ION CHANNEL THERMODYNAMICS STUDIED WITH TEMPERATURE JUMPS MEASURED AT THE CELL MEMBRANE

**DOI:** 10.1101/2022.07.11.499616

**Authors:** Carlos A Z Bassetto, Bernardo I Pinto, Ramon Latorre, Francisco Bezanilla

## Abstract

Perturbing the temperature of a system modifies its energy landscape thus providing a ubiquitous tool to understand biological processes. Here, we developed a framework to generate sudden temperature jumps (Tjumps) and sustained temperature steps (Tsteps) to study the temperature dependence of membrane proteins under voltage-clamp, while measuring the membrane temperature. Utilizing the melanin under the *Xenopus laevis* oocytes membrane as a photothermal transducer, we achieved short Tjumps up to 10 ºC in less than 1.5 ms and constant Tsteps for durations up to 150 ms. We followed the temperature at the membrane with submillisecond time resolution by measuring the time-course of membrane capacitance, which is linearly related to temperature. We applied Tjumps in Kir 1.1b, which reveals a highly temperature-sensitive blockage relief and characterized the effects of Tsteps on the temperature-sensitive channels TRPM8 and TRPV1. These newly developed approaches provide a general tool to study membrane proteins thermodynamics.

## Introduction

Temperature is an intensive physical property that results from the movement of atoms and molecules. As such, it affects from chemical reactions to biological processes. Temperature changes have been extensively used to study the properties of ion channels, and receptors, as a steady-state change of the bath temperature ^1,2,11,12,3–10^ or as a transient change by temperature jumps ^13–16^. Bath temperature changes are orders of magnitude slower than the gating kinetics of the channels, preventing the measurement of the system’s dynamics. Despite their appeal, fast temperature perturbation techniques have not been widely adopted to understand ion channel thermodynamics. Some challenges to fully implementing these methodologies arise from the technical difficulties of simultaneously achieving homogenous heating and reliably measuring the change in temperature at the membrane. Here, we introduce two complementary techniques that allow for studying ion channel thermodynamics by fast temperature perturbations. First, we used the endogenous melanin of *Xenopus leaves* oocytes as a photothermal transducer and a diode laser (447 nm) to raise the temperature of the oocyte membrane in a few milliseconds (up to 9 °C in 1.5 ms) in the same membrane region where the currents are measured under voltage-clamp. Second, we exploited the linear dependence of the membrane capacitance with temperature to track these fast temperature changes at the cell membrane. This capacitance-based temperature measurement (CTM) method allows us to measure the temperature at the membrane with sub-millisecond time resolution.

To demonstrate the suitability of these two methods in the study of ion channel thermodynamics, we use the inward rectifier potassium channel Kir1.1b and the non-selective cations channels transient receptor vanilloid 1 (TRPV1) and melastatin 8 (TRPM8). While TRPV1 is activated by heat ^3^, TRPM8 is a cold receptor ^17^. These three channels exhibit mild voltage dependence making them ideal candidates to test the effects of temperature using these newly developed techniques.

Our results show that in Kir1.1b, temperature changes induced an increase in the current through the open pore and relieved the rectification. The relief of rectification shows a high temperature dependence with an enthalpy change (ΔH) comparable to canonical temperature-sensitive channels. In both TRPV1 and TRPM8, the ionic current generated by Tstep during a voltage protocol shows a biphasic behavior, one associated with increasing the single channel conductance and the other with changes in the open probability of the channel. These results attest that we can effectively apply these tools to the study of ion channels. We expect that the combination of fast temperature changes and CTM can enrich our understanding of membrane protein thermodynamics.

## Results

### Temperature jumps (Tjumps) and Capacitance-based Temperature Measurement method (CTM)

To generate the fast temperature changes at the membrane (Tjumps), we placed a current-modulated diode laser on top of the Cut Open Voltage Clamp (COVC) setup^18^ (Fig. 1a and Extended Data Fig. 1a-b). We used a pair of microlens arrays to ensure a homogeneous illumination of the oocyte dome (400-500 μm of diameter) (Extended Data Fig. 1c-d). The illuminated area corresponds to the animal pole that contains melanosomes near the plasma membrane (∼1um) ^19^. The melanin heatups with the absorption of visible light, generating a temperature change at the oocyte membrane. To characterize these sudden temperature changes, we developed a new method based on membrane capacitance measurements using impedance, taking advantage of the fact that membrane capacitance changes with temperature ^20–22^.

**Fig. 1.**
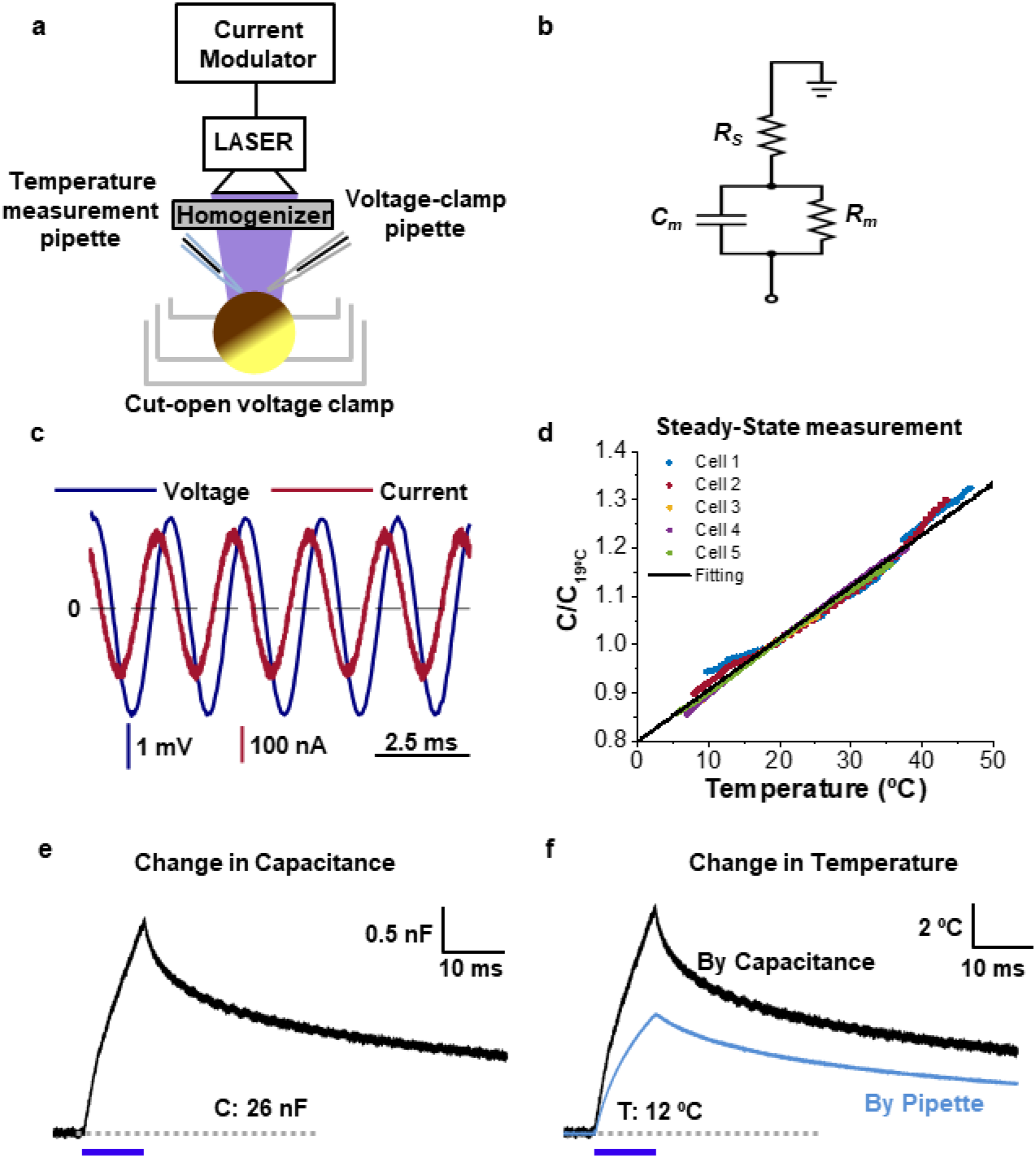
Tjumps and capacitance-based temperature measurement technique. **a**, Schematic representation of Tjumps setup. The voltage of oocyte dome was controlled using cut-open voltage clamp and at the same time it was illuminated using a homogenized laser beam. A calibrated pipette was positioned close to the membrane to measure the temperature. **b**, Equivalent circuit representation of the voltage clamped membrane. **c**, Applied sinusoidal voltage wave (blue) and elicited current (red). **d**, Relationship between normalized capacitance and temperature. The capacitance was normalized for each cell by dividing its value measured at 19°C. The line represents the regression line with slope of 1.070% ± 0.003 per 1 ºC. **e**, Capacitance change (obtained from the imaginary part of the impedance) during a laser pulse of 10 ms in duration. The duration of the laser pulse is indicated by the blue line. **f**, Temperature change obtained by CTM (black) and using a calibrated pipette placed near the oocyte membrane (light blue).

The impedance (*Z*) extends the concept of resistance into the frequency domain and is given by

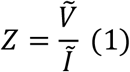

Where 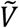 and *Ĩ* are the complex representation of a sinusoidal input voltage and the elicited current, respectively. Based on the equivalent circuit of the membrane (Fig. 1b), the impedance is given by:

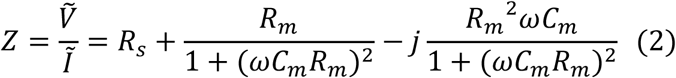

Where *R*_*s*_, *C*_*m*_, *R*_*m*_, *ω* and *j* are the series resistance, the membrane capacitance, the membrane resistance, the angular frequency of sinusoidal wave and the imaginary unit, respectively. From Eq. 2 and considering typical empirical values of *C*_*m*_ (∼ 18.2 nF) and *R*_*m*_ (∼ 1.17 Mω), we find that for frequencies larger than 300 Hz the term (*ωC*_*m*_*R*_*m*_)^2^ >> 1 (Extended Data Fig. 2). Then, the imaginary part of impedance becomes:

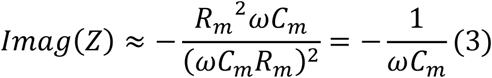

And the real part can be considered as:

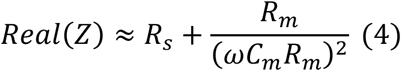

Thus, the impedance can be expressed as:

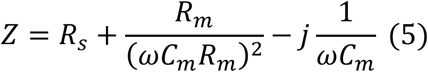

Eq. 5 demonstrates that we can obtain the membrane capacitance time course using the time course of the imaginary part of the impedance. To obtain the time course of capacitance with a high temporal resolution, we used the Hilbert transform, a powerful signal analysis tool that provides the imaginary component from a real-valued signal input ^23^. We, therefore, applied the Hilbert transform to the real-valued voltage and current signals to obtain their imaginary component, allowing us to compute *Z* (Eq. 1) and thus *C*_*m*_ (Eq. 3).

To determine the temperature dependence of membrane capacitance, we varied the bath temperature in the range of 4 - 45 °C. We took several measurements of the membrane capacitance after stabilization of the temperature at different values, monitored close to the oocyte dome using a thermocouple probe. The capacitance was estimated using a sinusoidal voltage command (Frequency 500Hz – amplitude 10 mV) while recording the resulting current (Fig. 1c). Since each cell has similar values for the total capacitance, we decided to normalize the capacitance by its value at 19 °C and found that capacitance is approximately linear within this temperature range and changes 1.070% ± 0.003 per 1 °C (Fig. 1d). This finding is consistent with previous calculations by Taylor using squid giant axons ^20^.

Finally, we use this linear relationship to convert changes in capacitance to changes in temperature, which is the foundation of our capacitance-based temperature measurement (CTM) method. We used the following procedure to calculate the changes in capacitance. First, we recorded the optocapacitive currents (Iop), elicited by Tjump, under conditions where there was no ionic conduction (Extended Data Fig. 3a - Methods). These optocapacitive currents arise due to a light-induced change in capacitance produced by the heating of the membrane through a photothermal effect ^21,22^. Low membrane conductance, which is obtained either in the absence or by preventing conduction of ion channels, provide high values of *R*_*m*_ (>1MΩ), which allows us to use the approximation in Eq. 3 for the frequencies used here (<1kHz). Next, we used a sinusoidal voltage wave to elicit currents while Tjump was applied (Isine + Iop, Extended Data Fig. 3b). This current is the combination of the sinusoidal current plus the optocapacitive current. Thus, to isolate only the sinusoidal component, we subtract Iop (Extended Data Fig. 3c). The subtracted current Isine can be used to calculate the imaginary component of the impedance according to Eq. 1 using the Hilbert transform. Finally, we used the imaginary part of the impedance to obtain the time course of *C*_*m*_ according to Eq. 3 (Fig. 1e). We validated this approximation by calculating *C*_*m*_ using three different frequencies and fitting Eq. 2 to the time course of *Z* (Extended Data Fig. 4a). We found a negligible difference between the fitting using Eq. 2 and the approximation using the Eq 3 for the tested frequencies (Extended Data Fig. 4b). Therefore, hereafter we use a single 500Hz frequency in all our experiments to determine *C*_*m*_.

Using the relationship found in Fig. 1d, we can now convert changes in capacitance to changes in temperature at the membrane, with an expected error of 0.03ºC per every 10º (Fig. 1f). To confirm the accuracy of the CTM method in measuring transient temperature changes at the membrane, we compared it with the changes in temperature using a calibrated glass pipette placed as close as possible to the oocyte membrane (Fig. 1a - Methods). The temperature changes calculated by CTM, and calibrated pipette have similar profiles; however, they differ in amplitude (Fig. 1f). This is expected since CTM provides the temperature change at the membrane compared to the pipette that is detecting the temperature changes micrometers away from the membrane. Therefore, CTM offers a straightforward, more accurate and probe-less way of measuring the temperature changes at the membrane where the proteins are embedded.

### Effects of Tjumps on ionic conductance and rectification of Kir

Next, we set to test the effect of Tjumps on ionic conductances of ion channels using the inward rectifier K^+^ channel Kir 1.1 isoform b ^24^. The voltage dependence of the Kir fammily channel arises from a fast internal block by magnesium and polyamines ^25^. Kir 1.1 is a weakly inward rectifying channel. To isolate the Kir ionic currents from other temperature induced currents, such as optocapacitive and leak currents, we used a linear subtraction procedure (Extended Data Fig. 6). This was implemented after blocking Kir currents by extracellular Ba^+2^ (Methods). When Tjumps were applied, an increase in ionic currents was observed, consistent with an increase in the K^+^ conductance (Fig. 2a). Taking the isochronal of the elicited current and knowing the temperature, it is possible to obtain I-Vs for several temperatures in a single experimental protocol (Fig. 2b). The direction and amplitude of the current follows the K^+^ reversal potential (V_rev_ ∼ -57mV). At voltages where V < V_rev_ Tjumps induces an inward current, whereas an outward current is observed when V > V_rev_. For V < V_rev_, the laser-induced currents exhibit a similar time course of the temperature change induced by the laser pulse (Fig. 2a). Since in these channels the rectification occurs at voltages larger than V_rev_, we interpreted the effects of Tjumps on these voltages as an increase in the single-channel conductance (increasing the diffusion rate of K^+^ ions through the pore of the channel). From the currents at -80 mV we estimated a Q_10_ of 1.6 ± 0.2 for the single-channel conductance (Eq 6 - Methods), consistent with values found for other Kir channels ^26,27^. At hyperpolarizing voltages, the time course of the currents follows the time course of temperature, however for large depolarizing voltages, we observed that the time course of the currents deviates from the temperature-time course, suggesting a secondary effect other than the increase in the single-channel conductance (Extended Data Fig. 5). Increasing temperature produces a rightward shift of the conductance vs voltage (G-V) curve indicating a relief of rectification (Fig. 2c). We then, calculated the voltage at which the rectification inhibits 50 % of the channels (Vr_1/2_ Eq. 8 - Methods). We obtained V_r1/2_ for the different G-Vs, plotted them against their respective temperature, and observed that V_r1/2_ shifts about 7 mV per ºC. The slope z of the G-Vs for the different temperatures shares the same value (1.25). From fittings of Vr_1/2_ with Eq. 9, we obtained the entropic (ΔS = -263 ± 10 cal/K.mol) and enthalpic (ΔH = -71.7 ± 2.9 kcal/mol) term of the rectification.

**Fig. 2.**
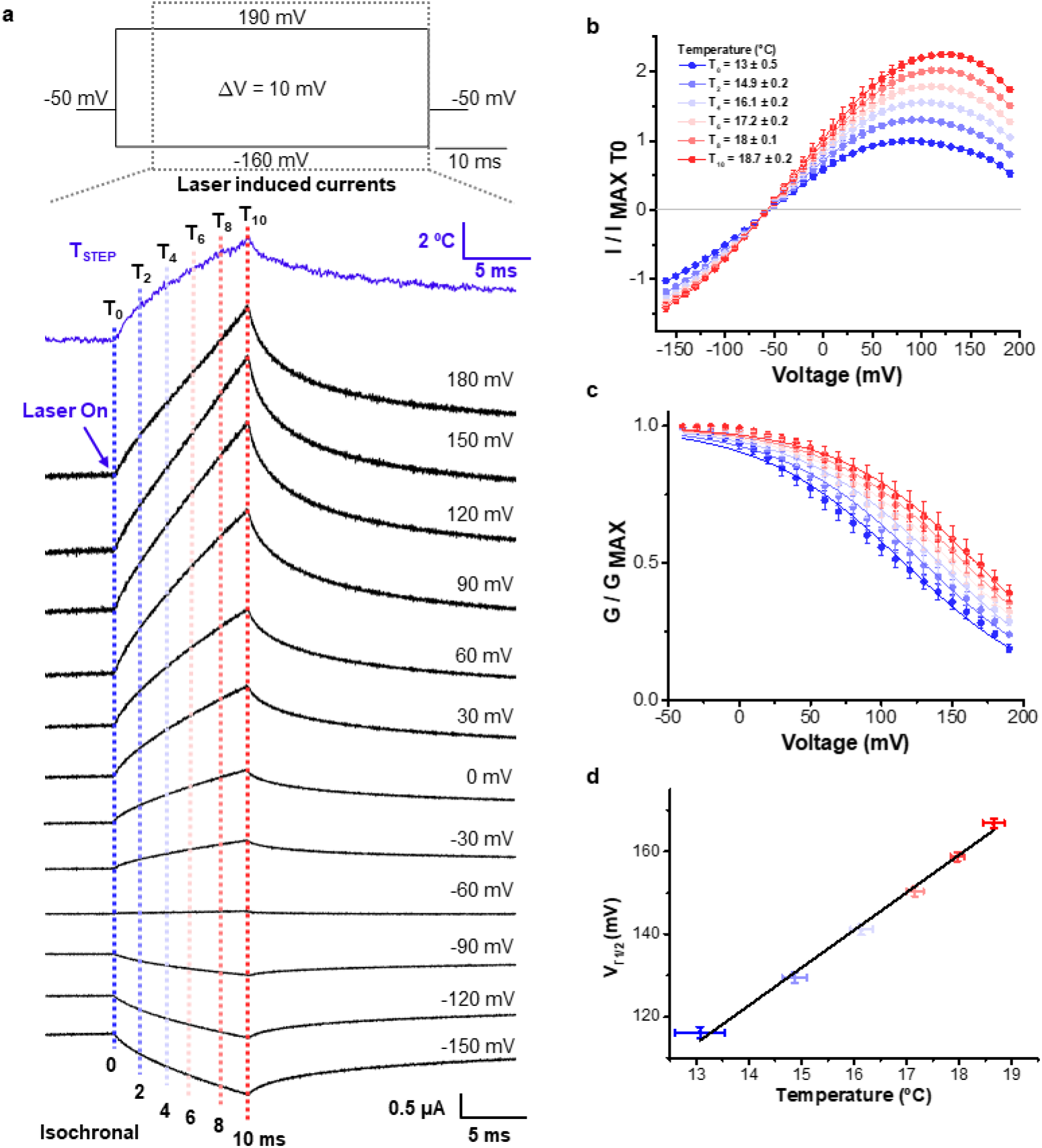
Effects of Tjumps on Kir ionic currents. **a**, Currents induced on Kir by 10ms Tjump at different voltages. Inset is the voltage protocol. The temperature change measured by CTM is shown in blue, the arow indicates when the Tjumps were started. The dashed lines indicate isochrones used to obtain the I-V curves. **b**, Normalized currents (I/ImaxT0) vs voltage for the different isochrones and their respective temperature (T0∼13 ºC) **c**, G-V relationship of Kir for different Temperatures. The G-V at different temperatures were fitted using Eq 8 with a shared slope (z = 1.25) for all the curves. **d**, Temperature-dependence of V_r1/2_ obtained from the G-V fits in C. The experimental values were fitted using Eq. 9 described in the methods section. The fitted values were ΔS = -263, ± 10 cal/K.mol and ΔH = -71.7 ± 2.9 kcal/mol. Error bars indicate standard deviation (*n* = 3).

### Temperature Steps

Next, we expanded the Tjumps technique to develop a sustained Temperature Step (Tstep) technique by modulating the durations of a train of laser pulses. As expected from a heat diffusion process, membrane heating is governed by the absorption of light by melanin, the distance between melanin and the membrane, the width of melanin layer, and the power of the laser. Cooling is a slower process governed by heat diffusion into the solution. Considering these characteristics of the heating and cooling of the membrane, we empirically determined a set of pulses to achieve a Tstep using pulse width modulation (PWM). We apply an initial pulse of 1-1.5ms followed by brief and repetitive pulses with a constant duration of 50-25μs but with a variable time between pulses (Fig. 3a - Methods).

**Fig. 3.**
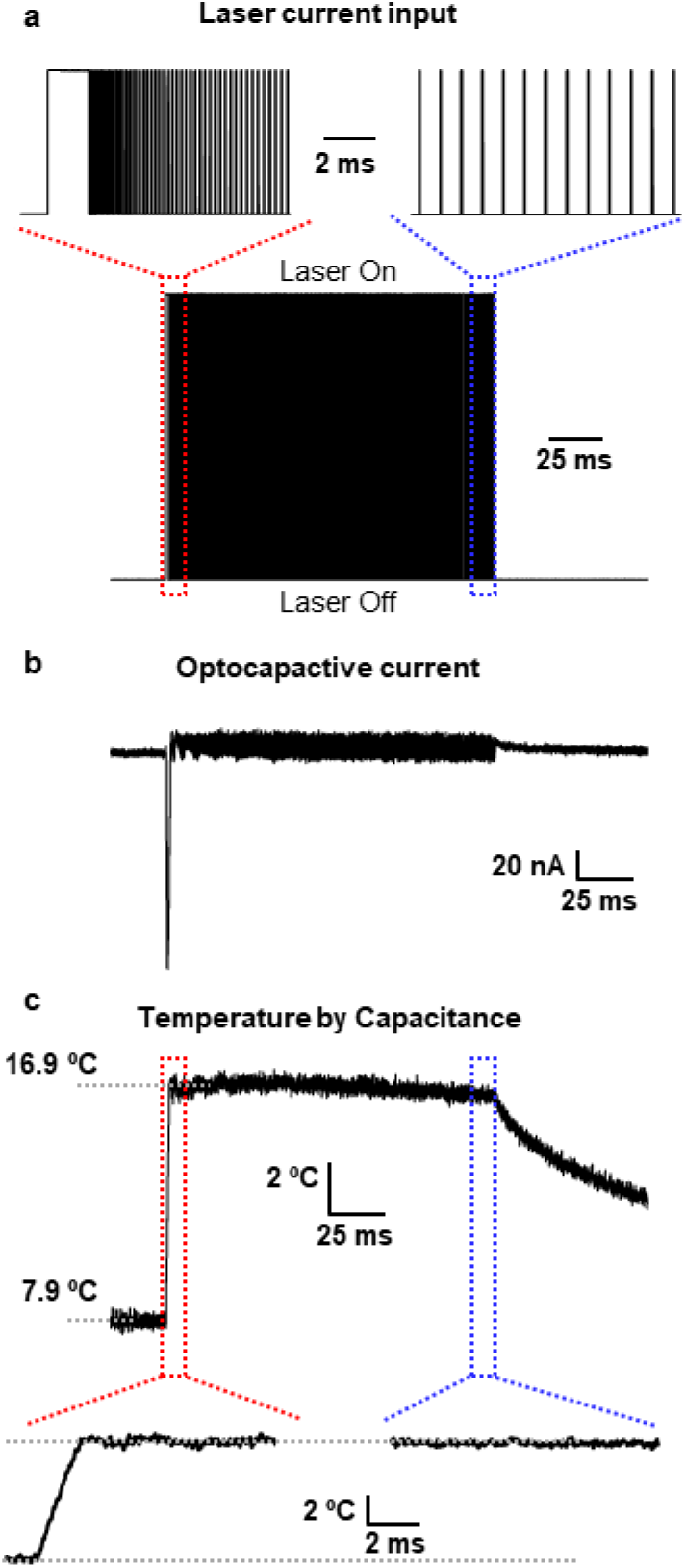
Using the PWM of the laser to build up Tsteps. **a**, Modulation waveform utilized to obtain a Tstep. Insets show an expanded time window to appreciate better the PWM pulses used. Red represents the beginning and blue for the end of the PWM pulses. **b**, Optocapacitive current elicited by Tstep at a holding potential of - 100mV. **c**, Change in temperature measured by CTM method. Insets are the expanded time window for the rising phase of temperature (red) and the Tstep end (blue).

This pulse protocol produces a large optocapacitive current during the rising temperature phase. Since the subsequent laser pulses maintain the temperature constant, they do not produce large optocapacitive currents (Fig. 3b). We used CTM to monitor the temperature changes, and it is possible to observe that the temperature increases by 9 ºC in 1.5 ms and keeps constant for 150 ms until the laser pulses are stopped (Fig. 3c).

#### Modulating thermoreceptors with Temperature steps

Next, we tested Tstep and CTM techniques to study the thermo transient receptor channels: TRPM8 and TRPV1. First, we tested the ability of Tsteps to replicate the effects of bath temperature changes. For TRPM8, we cooled the bath temperature to 9.5 ºC, and for TRPV1, we maintained the bath temperature at 21 ºC. The currents were elicited using a voltage protocol in the presence and absence of Tsteps (7ºC) for both channels (Fig. 4a-b). When Tsteps were applied, inhibition and activation of the elicited currents were observed for TRPM8 and TRPV1, respectively (Fig. 4a-b). For TRPM8 the temperature step reduced 40-50% of the maximum elicited current produced by the depolarizing pulse (Fig. 4c), whereas for TRPV1 it increased 250-350% (Fig. 4d). In agreement with the biophysical properties of these channels previously characterized ^3,5,6,11,28^ the effects of Tsteps on TRPM8 and TRPV1 were opposite. These results demonstrate that Tsteps can be used as an alternative to the steady-state bath solution temperature changes.

**Fig. 4.**
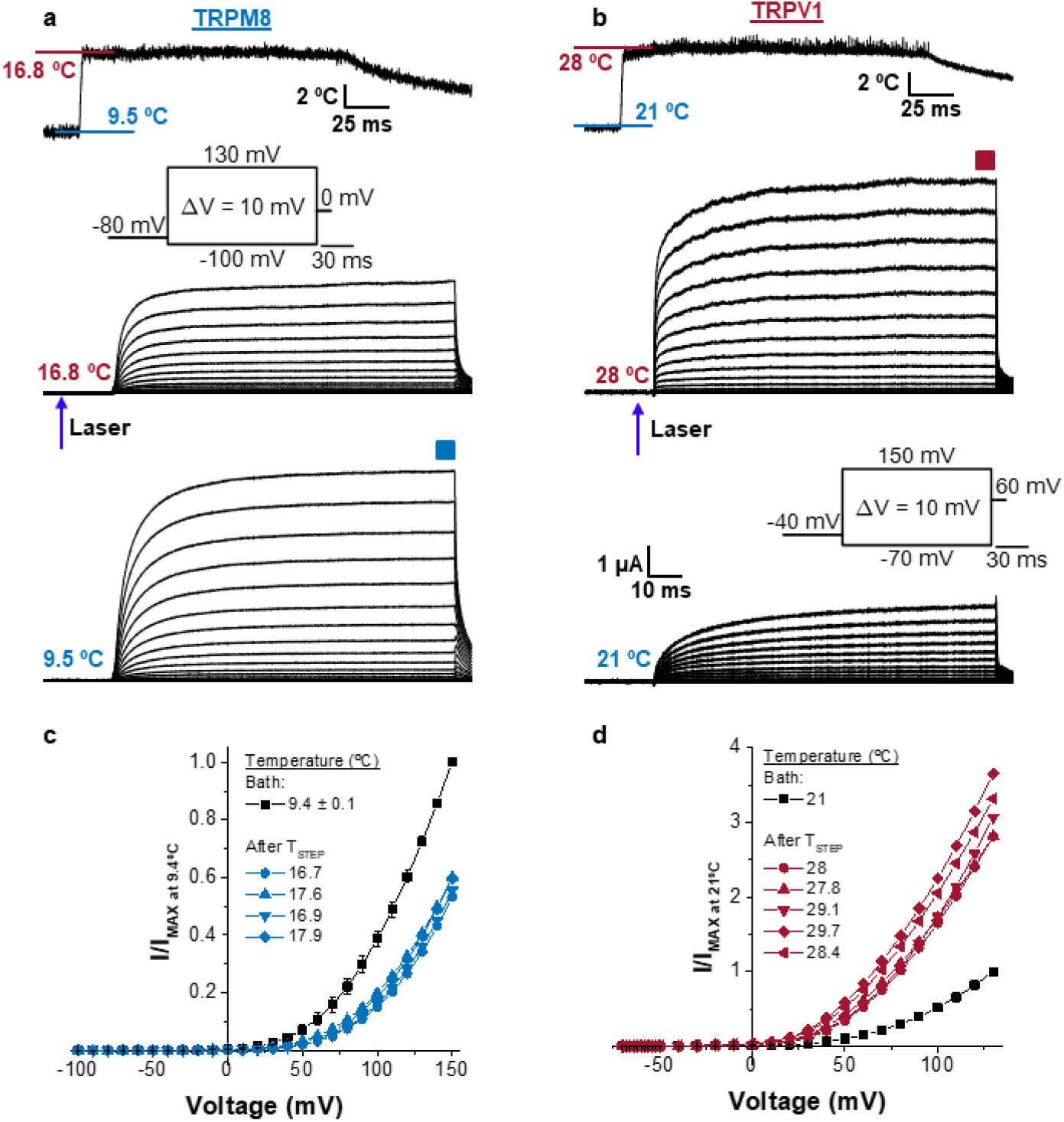
Effects of Tsteps on TRPM8 and TRPV1. **a**, and **b**, are, respectively, the representative current traces for TRPM8 and TRPV1. Top panel shows the Tstep for each case. Middle panel shows the current elicited under Tstep and bottom is the current without Tstep. Inset shows the voltage protocol used to elicit the currents. The arrow indicates the time where the temperature step was applied. The blue and the red square indicate the time at which the currents were taken to obtain the I-V curves. **c**, and **d**, are the I-V relationship for TRPM8 and TRPV1 ionic currents, respectively. The I-Vs were normalized by the maximum current recorded in absence of Tstep. Error bars indicate standard deviation for experiments in bath temperature condition (*n* = 4).

The technique developed here offers a new approach to perturb the system in a condition other than steady-state by applying a Tstep at any point during the activation process of the channels. To further explore this feature, we applied a Tstep in the middle of a voltage pulse on TRPM8 and TRPV1 (Fig. 5a-b). We observed the closure of TRPM8 and the opening of TRPV1 by Tsteps for all the voltages tested. Also, it is possible to observe that the currents exhibit a biphasic time course (Fig. 5c) with the Tstep. This biphasic effect is observed as a peak on the Tstep-induced currents in TRPM8 and an inflection point in TRPV1. The first phase is consistent with an increase of the single-channel conductance (Δγ), similar to Kir1.1b (Fig. 2). The second phase reflects the channel opening or closure (ΔP_o_) and starts about ∼ 200 μs after the onset of Tstep (Fig. 5c). The P_o_ changes rapidly until the temperature reaches steady state level, after this the Po continues changing although with slower kinetics. We interpret these results in light of the allosteric mechanism proposed for TRPM8 where the kinetics of deactivation have two-exponential decays, one with a fast time constant of 100-1000 us and a slower one with 2–10 ms time constant at 10 ºC ^28^. It is expected that the fast component of the current develops during the initial temperature change (1.5 ms). Thus, the currents observed during the steady state of the temperature reflects mostly the slow component.

**Fig. 5.**
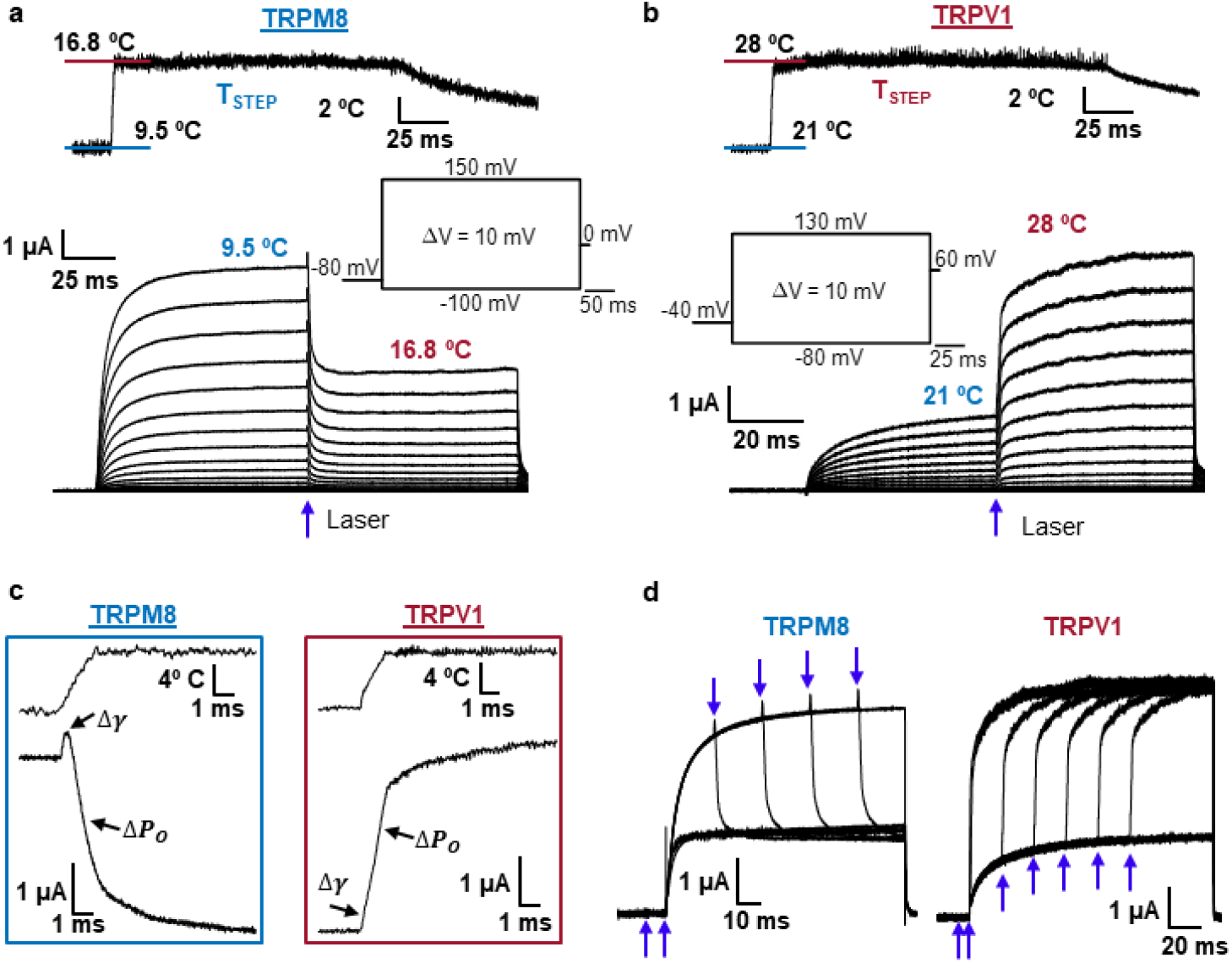
Time-dependence effects of Tstep on TRPM8 and TRPV1 ionic currents. **a**, and **b**, are ionic currents traces for TRPM8 and TRPV1, respectively. The temperature pulse is applied when the currents reach steady-state, as indicated by the arrow. Inset is the voltage protocol and Tstep profile. **c**, Biphasic behavior of currents elicited by Tstep for TRPM8 at 150 mV and TRPV1 at 120mV. The arrows indicate changes associated with Δγ and ΔP_O_. **d**, Tstep applied at different times during a voltage protocol for TRPM8 and TRV1. Arrows indicate different times where Tstep was applied.

The current can be up to threefold larger or twofold smaller than the ionic current obtained before the Tstep for TRPV1 and TRPM8, respectively (Fig. 5c). Using this technique, we can observe the kinetics of the channels responding only to temperature at a given kinetic point for the set voltage. Finally, we applied Tstep before and during a voltage step from -80 to 100 mV for TRPM8 and from -40 to 120 mV for TRPV1. The kinetics depend on the time when the Tstep was applied, but the steady-state current elicited by Tstep reached the same value irrespective of the time when the Tstep was applied for both channels (Fig. 5d).

## Discussion

Through the development of the CTM, we implemented Tjumps, and Tteps techniques to study ion channel thermodynamics. Our approach overcomes limitations in the measurement and control of membrane temperature and offers new insights into the function of ion channels. First, the CTM method allows us to measure the temperature directly at the membrane level using capacitance measurements and the known relationship between temperature and capacitance. This method relies on the Hilbert transform for fast capacitance measurements via impedance. The Hilbert transform provides a key advantage: time resolution is higher than the carrier wave frequency. We believe this approach can be employed in other settings where capacitance measurements are routinely used, such as the study of vesicular fusion ^29^, and the movement of gating charges ^30–32^. The CTM method provides a compelling advantage: an accurate readout of the temperature at the membrane level without needing of external probes. This mitigates problems that arise with the use of external devices to measure temperature, such as electrical noise, concerns about the positioning of the probe and the need for constant calibration in between experimental conditions. Second, we have developed a new approach to generate Tjumps and Tsteps at the cell membrane, which relies on light absorption by the melanin located on the animal pole of *X. laevis* oocyte.

Tjumps techniques have been utilized previously on nodes of Ranvier from *X. laevis* frogs ^33,34^, oocytes ^35^ and in cultured cells ^15,16,36,37^ using infrared light absorption by water or visible light absorption by graphite. However, we believe that the technique presented in this study offers some advantages. First, previously Tjumps were employed using the Two-Electrode voltage-clamp technique, where the whole oocyte is under voltage-clamp. In this configuration, the illuminated area is smaller than the area where the currents are recorded. The net recorded current comes from parts of the membrane that are under different temperatures. This does not happen in here because we used the COVC technique, and a homogeneous laser-focused on the oocyte dome under voltage-clamp and from where the currents are measured. Therefore, all the area under the voltage control is at the same temperature. Second, since our approach uses a blue laser instead of infrared, our method has three advantages. i) it produces the heat close to the cell membrane avoiding unintended heating of the solution ii) dissipates faster because the heating is localized and iii) penetrates deeper into the solution than middle infrared.

When obtaining the temperature dependence of an ion channel, it is necessary to increase the bath temperature, wait for a few minutes for the temperature to reach the desired value and then record the currents. Since this process can take a long time, problems like rundown may arise during a set of experiments. Tjumps and Tstep techniques offer a better option, as it is possible to get the temperature dependence of current in a single long pulse that can be set to reach different temperatures (Fig. 2, 4, and 5). Therefore, we recorded pairs of currents for the same voltage protocol being one in presence and the other in the absence of Tstep. Using this pattern of recordings, we can always check whether the laser pulse is not damaging the membrane, avoiding the rundown of currents that usually appear during the long times required when the recordings are done by steady-state controlled bath temperature.

When used in combination, Tsteps and CTM provide a powerful tool to modify the energy landscape that defines the gating of ion channels without the need for additional temperature probes. These features are appealing because one can assess the temperature dependence of thermodynamic processes. To show the applicability of these methods, we have used Kir1.1b, TRPM8 and TRPV1 channels. Our observations reproduce results previously reported changing the temperature of the bath, but also revealed new striking effects of temperature on these channels: i) the biphasic time course of currents elicited by Tstep on TRPV1 and TRPM8. These results imply that the temperature deactivation process in TRPM8 follows a double exponential time course as previously reported^29^. We found the same double exponential time course for the activation in the case of TRPV1 suggesting that similar allosteric gating kinetic models^29^ may explain the coupling between voltage and temperature sensor in TRPM8 and TRPV1; ii) the large temperature dependence of rectification in the Kir1.1b channel. The Kir1.1b channel rectification has a large enthalpic component, comparable to some thermo TRPs^38^. The rectification of Kir channels arises from the block of the pore by cytoplasmic polyamines and magnesium ^25,39,40^. The block of the channel would produce a decrease in entropy of the blocker molecule. For the blockage to be stable this entropy change needs to be offset by an enthalpy change. It is possible that the interaction between the positive charges of the blocker with negatively charged amino acids located in the internal vestibule provide this large enthalpic changes and thus give rise to the temperature dependence observed.

We believe that Tstep is a powerful tool to study the landscape of energy as the ion channel evolves from one kinetic state to another. The voltage-clamp technique allows to impose a membrane voltage at constant temperature and then change it to the desired value to obtain the voltage-dependence of the kinetics of an ion channel process. Similarly, the Tstep technique now offers a way to get the temperature-dependence of those transitions between kinetic states under constant voltage. For example, in Fig. 5 we showed that Tstep can be applied before, after, or while the ionic currents of the TRPM8 and TRPV1 were evolving, thus providing insights into the temperature dependence of the transitions between close to open and open to close of the channels.

We have illustrated the present method with an inward rectifier and two thermoreceptor channels, but we envision that the methods and approaches described in this work can be applied to understand the general gating mechanisms of different families of ion channels, or any membrane protein that its function may be studied with voltage clamp, such as many transporters and pumps, because temperature affects their landscape of energy. Furthermore, the approach used to generate Tstep can be extended to other preparations, *e*.*g*., in combination with the patch-clamp technique. This could be achieved by injecting melanin via the patch pipette or decorating the cells with gold nanoparticles^41^. It would require the use of small enough cells so that the temperature gradients from the illuminated to the non-illuminated side are not too different. This would allow the application of Tstep to study isolated neurons and cells, enriching the knowledge of the temperature effects on those preparations.

## Materials and Methods

### Channels expression in *Xenopus* oocytes

*Xenopus laevis* oocytes were surgically harvested following experimental protocols #71475 approved by the University of Chicago Institutional Animal Care and Use Committee (IACUC). The follicular membrane was digested by collagenase 2 mg/ml supplemented with bovine serum albumin (BSA) 1mg/ml. Oocytes were kept at 12 or 18 ºC in SOS solution containing (in mM) 96 NaCl, 2 KCl, 1 MgCl_2_, 1.8 CaCl_2_, 10 HEPES, pH 7.4 (NaOH) supplemented with gentamicin (50 mg/ml). They were injected after 6 - 24 hr of harvesting, with 5-50 ng of cRNA diluted in 50 nl of RNAse-free water and incubated for 1-4 days prior to recording. We used the rat renal inward rectifier Kir1.1b (ROMK2) cloned in pSPORT vector (kindly provided by Dr. Henry Sackin), rat TRPM8 and rat TRPV1 channels cloned into pBSTA vector. cRNAs were transcribed using the mMESSAGE mMACHINE T7kit (Life Technologies, Carlsbad, CA) and linearized with NotI-HF (New England Biolabs, Ipswich, MA). cDNAs were sequenced to attest to the right sequence.

### Electrophysiology

Ionic currents were recorded from oocytes using the cut-open voltage-clamp method ^42^. Voltagesensing pipettes were pulled using a horizontal puller (P-87 Model, Sutter Instruments, Novato, CA), and the resistance ranged between 0.2-0.5 MΩ. Data were filtered online at 20–50 kHz using a built-in low-pass four-pole Bessel filter in the voltage clamp amplifier (CA-1B, Dagan Corporation, Minneapolis, MN, USA) sampled at 1 MHz, digitized at 16-bits and digitally filtered at Nyquist frequency (USB-1604; Measurement Computing, Norton, MA). The voltage command and the current elicited were filtered using the same frequency. The current traces for TRPs and the temperature traces obtained by CTM were offline filtered at 10KHz. An in-house software was used to acquire and analyze the data (GPatch64MC). The chamber temperature was measured by a thermocouple and controlled through a negative feedback loop using a Peltier cooler. Transient capacitive currents were subtracted from the recorded currents by a dedicated circuit. For ionic current measurements, the external solution was composed of (mM): KOH 12, CaOH_2_ 2, HEPES 10, EDTA 0.1, N-methyl-D-glucamine (NMDG) 108, for Kir1.1 and TRPM8; and NMDG 120, MgOH_2_ 2, HEPES 10 for TRPV1. In all cases, the internal solution was composed of (mM): KOH 120, EGTA 2, HEPES 10. All the solutions were adjusted to pH 7.4 with methanosulfonic acid. All chemicals used were purchased from Sigma-Aldrich (St. Louis, MO). To block Kir1.1b currents we added BaCl_2_ to the top and guard chambers of the COVC to a final concentration of ∼1mM.

### Data Analysis

To estimate the temperature dependence of the single-channel conductance for Kir, we obtained the Van’t Hoff plot for the currents at -80 mV, where there is no rectification. The currents during the Tjump were normalized by dividing them by the mean current before the Tjump. The logarithm of this current ratio was plotted against *1/T*. From the slope of the Van’t Hoff plot we obtained the *ΔH* and hence the *Q*_*10*_ for the currents using the equation ^43^:

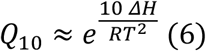

The I-V curves were transformed into conductance (*G*) by using the equation below:

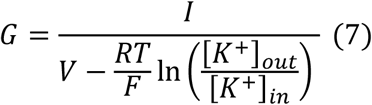

where *I* is the K^+^ current activated by membrane voltage *V, R* is the gas constant, *T* is the temperature in Kelvin, *F* is the Faraday constant, and [*K*]_*in*_ and [*K*]_*out*_ are the intracellular and extracellular K^+^ concentrations, respectively.

The individual K^+^ conductances were then normalized, averaged, and plotted against *V* to build conductance-voltage (G-V) curves. The conductance data were fitted using a two-state model given by the following equation

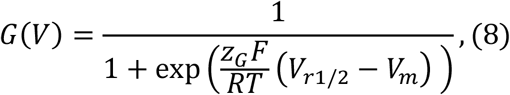

where *z* is the apparent charge of the rectification process expressed in units of elementary charge (*e*_0_) and *V*_*r*1/2_ is the voltage at which rectification inhibits 50% of the channels.

The Vr_1/2_ values calculated using for the different temperatures were fitted using the linear relation _44_:

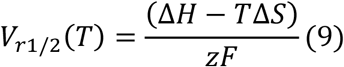

Data analysis, graphs, and modeling were performed using programs written in-house (Analysis), MATLAB 2022a (The MathWorks, Inc, Natick, MA), MATHEMATICA 12 (Wolfram, Champaign, IL), Origin 9.0 (Origin Lab Corporation, Northampton, MA).

### Tsteps set up

A 3.5 W 447nm diode laser (Osram PLPT9 450D_E A01) was placed on top of the recording chamber and aligned with the oocyte dome (Fig. 1a). The beam was collimated using an aspheric lens (A230TM-A,Thorlabs) followed by a pair of microlens arrays (MLA300-14AR-M, Thorlabs Inc. Newton, New Jersey) to ensure homogenization of the beam ^45^, and focused into the oocyte dome using another aspheric lens (ACL2520U-A,Thorlabs Inc. Newton, New Jersey). (Extended Data Fig.1a-b). A specially designed high current, modulated power supply was used to achieve rapid turn-on of the laser. The laser was gated by a TTL pulse from the same DAQ used to control the voltage clamp or by an arbitrary wave generator (4075B, B&K Precision corp., Yorba Linda, CA).

To generate Tsteps, we apply an initial long pulse of 1-1.5ms followed by brief repetitive pulses of constant duration of 50-25μs with a variable time between pulses (Fig. 3a). The time in between laser pulses increases with each repetition following the equation:

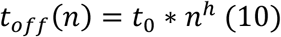

Where n, t_off_ and t_0_ are pulse number, the time in between laser pulses and the initial off time, respectively. The h factor is an empirically adjusted exponent that determines how fast the time between pulses increases and is usually between 0.7 and 0.9.

### Subtraction of linear components

We used a linear subtraction procedure to isolate the effects of Tjumps and Tsteps on optocapacitive currents ^21^ and leak currents. In the case of TRPs we used voltages in the range where no ionic currents are present, while for Kir we used external barium to block inward ionic currents. These currents were fitted using a linear equation and subtracted by extrapolation from the traces where ionic currents are present. For example, Extended Data Fig. 6a-b show a representative current recorded from an uninjected oocyte at different voltages in presence of Tjump. The subtracted current is shown in Extended Data Fig. 6c. The colored dashed lines are 4 isochrones taken to exemplify the subtraction procedure. Extended Data Fig. 6d shows the I-Vs from four isochrones taken at different times for the experimental data (I_exp_) with their respective linear fittings (I_fitted_) and the results after subtraction (I_exp_ - I_fitted_).

### Resolution of the temperature measurements using CTM

We used Eq. 1 to calculate the impedance, dividing the voltage command by the current. As stated in the results section, it is necessary to subtract the optocapacitive current to calculate the change in capacitance. All the division and the subtraction procedure ended up building noise in the analysis. To decrease it, we recorded 100 times the optocapacitive current and 100 times the current elicited by the sinusoidal voltage wave. We averaged the optocapacitive current to reduce the noise, and then the averaged optocapacitive current was used to subtract each of the sinusoidal currents resulting in 100 subtracted traces. They were processed using the Hilbert transform to get the capacitance change. Finally, the final 100 traces of capacitance were average to get the change in capacitance and afterwards converted into changes in temperature. Even though all the efforts to decrease the noise, the final averaged trace of the temperature still contained some noise (Extended Data Fig. 7). Therefore, we used a 10KHz *offline* Bessel-like filter to obtain the final temperature-time course. We achieved a resolution of ∼0.5 ºC peak-to-peak (Extended Data Fig. 7).

### Temperature measurement using a calibrated pipette

The temperature measurement was based on the method developed by Yao and colleagues^15^. Briefly, pipettes with a resistance of 2–5 MΩ were filled with solution matching the extracellular recording buffer containing (in mM) 120 KCl, 5 HEPES, and 2 CaCl_2_, set to pH 7.4. The pipette tip was positioned in the bath above and as close as possible to the oocyte dome. Pipette resistance was monitored using an in-house voltage-clamp amplifier. A resistance-temperature calibration curve was obtained changing the bath temperature while simultaneously recording pipette resistance and the bath temperature with a thermocouple. A linear relationship was obtained using an Arrhenius plot, and this calibration curve was used to convert resistance changes measured during the application of the Tsteps into temperature (Fig. 1f).

## Acknowledgments

CB and BP agree that both authors should be the first author and the order of appearance was determined by coin toss. We would like to thank Dr. Sara T Granados for her help in designing Fig. 1a.

## Funding

National Institutes of Health Award R01GM030376 (FB, RL)

Fondo Nacional de Desarrollo Cientıfico y Tecnologico (FONDECYT). Regular Grant Number 190203 (RL)

PEW Latin American Fellow 2019 (BP)

The Centro Interdisciplinario de Neurociencias de Valparaiso (CINV) is supported by the Iniciativa Cientıfica Milenio - Agencia Nacional de Investigacion y Desarrollo (ICM-ANID), Project P09-022-F.

## Author contributions

Performed and analyzed experiments: CB, BP

Interpreted results: CB, BP, RL, FB

Conceptualization: CB, BP, RL, FB

Writing: CB, BP, RL, FB

Supervision: RL, FB

## Data and materials availability

All data presented in the paper will be made available to readers upon request. The codes and exemplary data will be deposited on Github.

