## Supplementary material for "ION CHANNEL THERMODYNAMICS STUDIED WITH TEMPERATURE JUMPS MEASURED AT THE CELL MEMBRANE": Su


**This PDF file includes:**

Extended data Figure 1 to 7

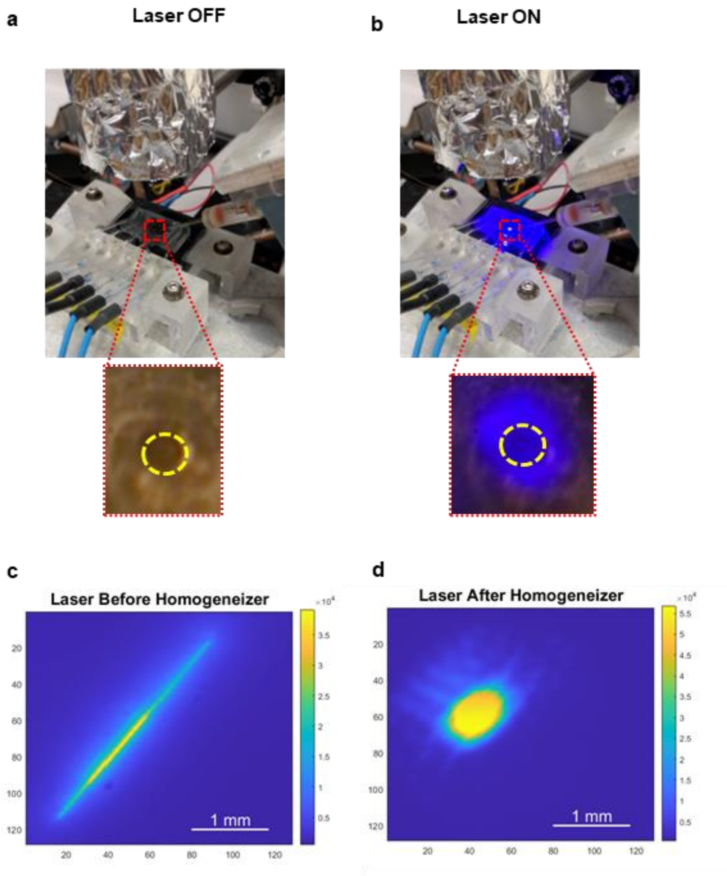


**Extended Data Fig. 1. Setup and laser light homogenization. a,** Picture of the COVC with the laser coupled to it. The laser is cover in thin foil to prevent electrical interference. **b,** same as **a**, but with laser on. Insets show the oocyte dome under voltage control (yellow circle) and the illuminated area. **c,** and **d,** are the laser beam intensity profile before and after homogenization, respectively. The calibration bar shown in the right of **c** and **d** indicates Arbitrary Intensity Units scale.

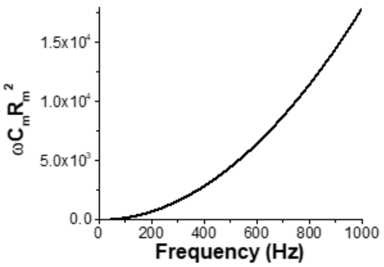


**Extended Data Fig. 2. Validation of the approximation shown in Eq 3.** Plot of $(\boldsymbol{\omega C_m R_m})^{\mathbf{2}}$ versus frequency. Graph generated with $C_m$ (18.2 nF) and $R_m$ (1.17 MΩ) taken from typical experimental values.

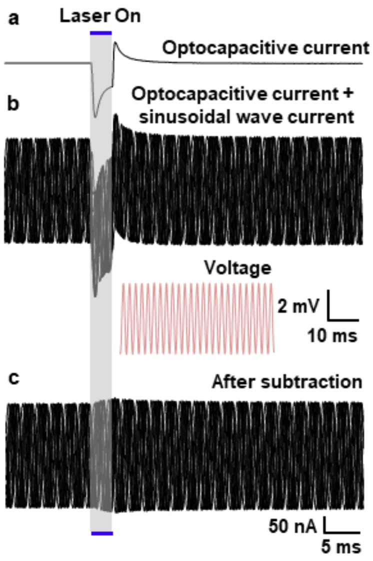


**Extended Data Fig. 3. Calculating the Impedance (Z) to obtain capacitance changes by Tstep. a,** Optocapacitive current (Iop) induced by Tjump. **b,** Overlapping of 10 current traces induced by sinusoidal voltage waves and Tsteps (Isine + Iop). The voltage wave shown in the inset (red) has an amplitude and frequency of 4mV and 500Hz, respectively. **c,** Current (Isine) obtained after subtraction of **b** from **a**. The holding potential was -80mV. Blue traces indicate the duration of the laser pulse.

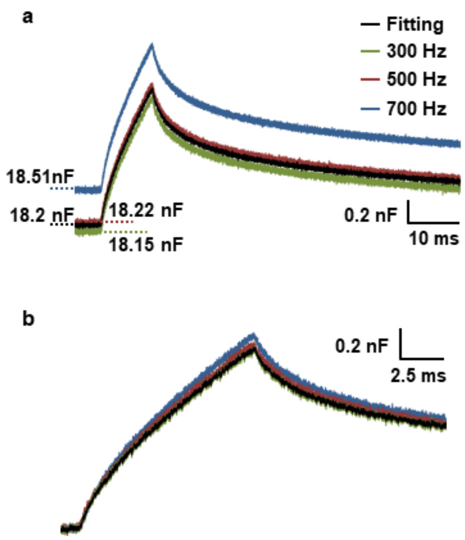


**Extended Data Fig. 4. Capacitance measurement validation. a,** Measurement of the change in capacitance using the fitting of the Z to Eq. 2 (black). The green (300 Hz), red (500 Hz), and blue (700 Hz) traces are the capacitance calculated using the approximation shown in Eq. 3. **b,** Superposition of capacitance traces shown in **a**. Note that the approximation used in Eq. 3 is in good agreement with the fitted data.

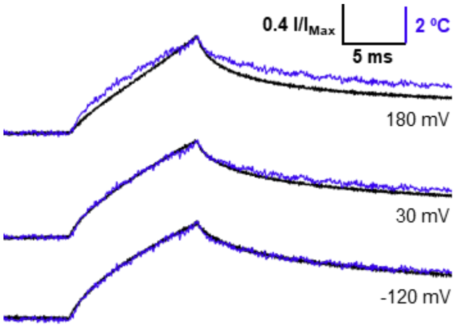


**Extended Data Fig. 5. Comparison between the time course of Temperature and normalized ionic Kir1.1b currents induced by Tjumps.** Temperature time course is shown in blue and elicited current by Tjump is in black. The currents were normalized by their relative peak amplitude. Note that due to this normalization, the current at -120 mV is flipped. Normalized currents and temperature are from the traces shown in Fig. 2a.

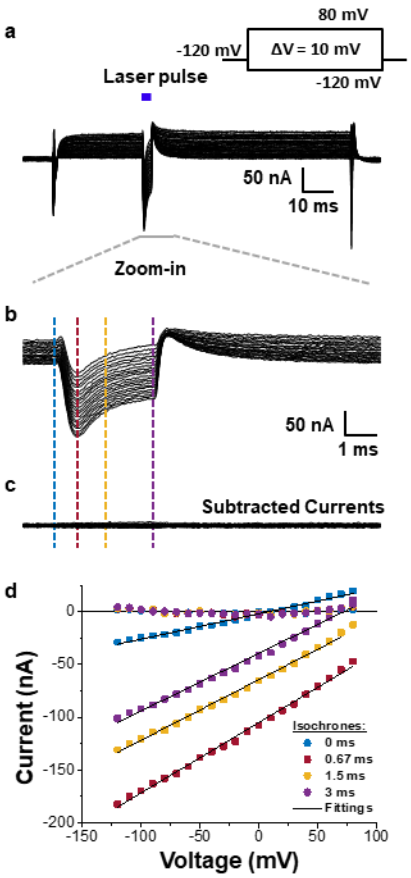


**Extended Data Fig. 6. Example of the subtraction procedure used to remove the linear components of the current and isolate the effects of Tjumps and Tsteps. a,** Ionic currents elicited by Tjump from uninjected oocytes blue line represents the duration of the laser pulse. Inset is the voltage protocol. **b,** Expanded time window of the elicited current shown in **a** for better appreciation of the effects of Tjumps. **c,** Ionic currents from **b** after the subtraction procedure currents. Dashed lines are isochrones to build the I-V curves. **d,** I-V curves from different times for subtracted (circles) and unsubtracted (squares) currents. The colors correspond to the isochrones in **b** and **c**. The solid line is the linear fitting used for the subtraction.

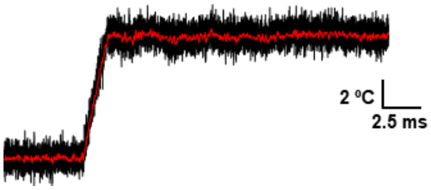


**Extended Data Fig. 7. Resolution of temperature measurements using CTM.** Temperature traces without *offline* filter (black) and *offlin*
